# A Mechanistic Reinterpretation of Fast Inactivation in Voltage-Gated Na^+^ Channels

**DOI:** 10.1101/2023.04.27.538555

**Authors:** Yichen Liu, Carlos AZ Bassetto, Bernardo I Pinto, Francisco Bezanilla

## Abstract

Fast Inactivation in voltage-gated Na^+^ channels plays essential roles in numerous physiological functions. The canonical hinged-lid model has long predicted that a hydrophobic motif in the DIII-DIV linker (IFM) acts as the gating particle that occludes the permeation pathway during fast inactivation. However, the fact that the IFM motif is located far from the pore in recent high-resolution structures of Nav^+^ channels contradicts this *status quo* model. The precise molecular determinants of fast inactivation gate once again, become an open question. Here, we provide a mechanistic reinterpretation of fast inactivation based on ionic and gating current data. In Nav1.4 the actual *inactivation gate* is comprised of two hydrophobic rings at the bottom of S6. These function *in series* and closing once the IFM motif binds. Reducing the volume of the sidechain in both rings led to a partially conductive inactivated state. Our experiments also point to a previously overlooked coupling pathway between the bottom of S6 and the selectivity filter.

## Introduction

Voltage-gated sodium (Nav) channels function as the main drivers of the depolarization phase of action potentials. As such, they are ubiquitously found in a wide range of excitable tissues, playing fundamental roles in many vital physiological functions, such as skeletal muscle contraction, heart beating, and neuronal impulse generation and propagation (1). Consisting of one long protein chain, mammalian Nav channels are arranged into four similar, yet not identical domains (DI to DIV). Each domain, formed by 6 transmembrane helixes (S1 to S6), is then organized pseudo-symmetrically into the <u>v</u>oltage-<u>s</u>ensing sub<u>d</u>omain (VSD, S1 to S4) and the <u>p</u>ore-forming sub<u>d</u>omain (PD, S5 toS6) (2). Milliseconds after activation, Nav channels rapidly enter into a non-conductive state, the fast inactivated state, and only recover upon repolarization (3–6). Fast inactivation serves as a negative feedback switch, responsible for the attenuation of inward current during action potential. Subtle abnormalities in fast inactivation can lead to severe pathological states (7–9). These diseases, though sharing similar etiology, can manifest in many different types of tissues and show a wide range of symptoms due to the prevalence and diversity of Nav channels (10). For instance, skeletal muscle hyperkalemic periodic paralysis (11, 12), long QT syndrome in the heart (13) and inherited epilepsy in brain (14) can all arise from abnormal Nav fast inactivation.

While examining the ionic and gating current from Nav channels in the giant squid axon, Armstrong and Bezanilla formulated the “ball and chain” model (15, 16), where an positively charge “inactivation ball”, tethered by a flexible “chain” to the intracellular domain of the channel, directly blocks the permeation pathway. The “ball and chain was thought to be part of an autoinhibitory process on the intracellular face of the channel, since fast inactivation is keenly susceptible to proteolytic action in internally perfused axons(17). Site directed mutagenesis experiments led to the identification of a triad of hydrophobic residues (Isoleucine-<u>Ph</u>enylalanine-<u>M</u>ethionine, the IFM motif) in the linker between DIII and DIV that is necessary for fast inactivation (18). Modifying the original “ball and chain” model, West *et al*. proposed a “hinged-lid” model, where the IFM motif was predicted to be the inactivation particle that physically blocks the pore during fast inactivation, acting as the inactivation gate. Many reports have subsequently supported this model, and the “hinged-lid” model was essentially accepted as the canonical model for fast inactivation in Nav channels (19, 20, 20, 21) until the advent of the cryoEM “resolution revolution”.

Upon the first mammalian Nav channel structure (22) it became clear that the IFM motif, though docking into a hydrophobic pocket, resided far away from the pore in the putative inactivated state to block the permeation path in contrast to the predictions of the canonical “hinged-lid” model. Since then, numerous structural reports of different isoforms of Nav channels all confirm this original observation (23–29). And while the IFM motif does not seem to serve as the fast inactivation gate, it is clearly necessary for fast inactivation process (18). As a result, the precise molecular determinants of fast inactivation gate have, once again, become an open question.

Here, we took advantage of the available structural data and tested directly the hypothesis suggesting that, rather than the IFM motif, fast inactivation in Navs is associated with large residues at the intracellular end of pore-forming helixes S6. Combining ionic and gating current measurements, we demonstrate and characterize a S6-located double ring of hydrophobic residues in DIII and DIV. Through site directed mutagenesis, we show that the fast inactivation gate is formed by two layers of bulky, hydrophobic residues, whose effectiveness as inactivation gate is dependent on side chain volume. Once reduced, smaller side chain substitutions lead to a leaky inactivated state. Equivalent residues in DII, on the other hand, seem to be involved in the coupling between VSD activation and PD opening instead. Surprisingly, our experiments point to a previously overlooked coupling pathway between the bottom of S6 and the selectivity filter (SF). Altogether, our results identify a new candidate and mechanism for the fast inactivation gate in Nav channels, highlighting importance of the intracellular (C-terminal) end of S6 for both activation and fast inactivation.

## Results

### Structural analysis reveals a two-tier hydrophobic barrier at the bottom of S6

Starting with the observation that the IFM motif is located far from the pore in the putative inactivated state (Fig 1A, B) (28, 30, 32), we reasoned that large, hydrophobic residues residing at the intracellular end of S6 might play a dual role participating both in activation and fast inactivation gating. Close examination of the electron density maps in the available Nav channel structures, shows that in most cases, non-protein densities can be spotted inside the pore, spanning across the bundle crossing region into the inner cavity (22, 32, 33). These molecules are sometimes unidentified however in many cases assigned to be detergent molecules, a fact that clouds any interpretation regarding their functional state. However, given that the structures are determined at 0mV and the VSDs are in the “up” conformation, it is reasonable to assume that the pore should resemble a fast inactivated state. Here we have focused our analyses on two structures that do not present such non-protein densities in the pore: NavPas (PDB:6A95 (30)) and Nav1.7 M11 (PDB:7XVF (28), referred to as Nav1.7 from now on) (Fig.1 A, B). We calculated the pore profile in NavPas and Nav1.7 (HOLE) as the radius along the permeation pathway (graphs in Fig.1 A, B), this profile led to easy identification of the narrowest part of the pore in each case. While NavPas and Nav1.7 showed very similar pore profiles at the intracellular end, the stretch along the pore that makes its narrowest part, was much larger in NavPas and Nav1.7 when compared to structures with non-proteinaceous densities inside the pore, as in the case of Nav1.5 structure (Supplementary Figure1). The narrowest region in NavPas and Nav1.7 was composed by two layers of hydrophobic residues organized in a diamond shape facing towards the pore at the bottom of S6, seen as two minima in the radius profiles (Fig. 1A, B), located one α helix turn away from each other. An early indication of the feasibility of our hypothesis is related to the fact that the 8 residues (two from each domain) identified in NavPas and Nav1.7, are highly conserved among different Nav isoforms, specially to rNav1.4 used in this study (Fig. 1C, Supplementary Figure2). All the residues identified were hydrophobic and 7 of them were also bulky. Based on our structural analysis, we hypothesize that these S6 residues act as hydrophobic barriers and potentially form the fast inactivation gate in Nav channels.

**Figure 1:**
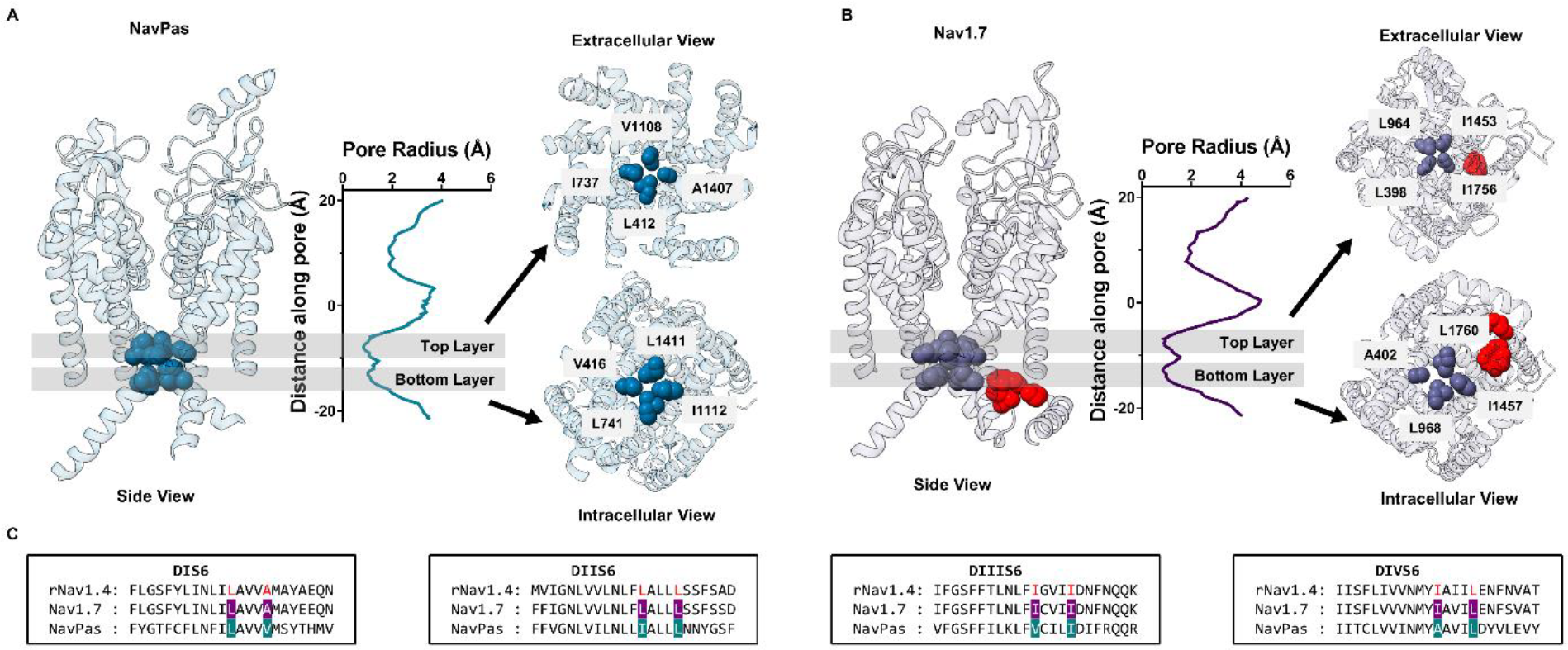
Pore radius profile of NavPas and Nav1.7. A) Structure of NavPas (PDB:6A95 - (30)). B) Structure of Nav1.7 M11 (PDB:7XVF - (28). The IFM motif is highlighted in red. The pore radius calculated using HOLE software (31). NavPas and Nav1.7 structures show similar pore profile at the bottom of S6, where two layers of hydrophobic barrier are identified (shadowed regions). Identified top (extracellular view) and bottom (intracellular view) hydrophobic layers in NavPas and Nav1.7 are shown. In NavPas, the top layer is formed by residues L412 (DI), I737 (DII), V1108 (DIII), A1407 (DIV) and the bottom layer by V416 (DI), L741 (DII), I1112 (DIII), L1411 (DIV). In Nav1.7, the top layer is formed by residues L390 (DI), L964 (DII), I1453 (DIII), I1756 (DIV) and the bottom layer by A402 (DI), L968 (DII), I1457 (DIII), L1760 (DIV). C) Sequence alignments of S6 region among rNav1.4, Nav1.7 and NavPa. The residues identified to form the narrowest part of the channel were highlighted in light blue for NavPas and dark blue for Nav1.7, and they are colored in red for rNav1.4.

### Double alanine mutation in DIII S6 produces a conductive “leaky” inactivated state

After identifying residues that could potentially form the fast inactivation gate at S6, we predicted that by reducing the volume of the involved side chains, we could widen the two-tier barrier in S6 and partially open the fast inactivation gate. We first tested DIII and DIV due to their known role in fast inactivation, and the fact that the IFM motif is spatially closer to these two domains (Fig. 1B) (34–36). In DIII, isoleucine I1284 and I1288 were identified based on the sequence alignment (Fig. 2 A). Single alanine mutations on either position (I1284A or I1288A) yielded currents similar to wild type (WT) channel (Fig 2A). When both residues were mutated to alanine simultaneously (mutant DIIIAA), significant steady state current was observed (Fig 2A, bottom traces). At +60mV this steady state current corresponds to around 20% of the peak current which clearly differs from the negligible amounts (less than 3%) in the WT as well as in the I1284A and I1288A mutants (Fig 2B). A small right shift, around 10mV, can be seen in the voltage dependence of the fast inactivation (h-infinity curve) in I1284A, I1288A and DIIIAA (Fig. 2C, Supplementary Table 3). The steady state current was present exclusively in the double mutant, which cannot be accounted for by a combination of the effects of the two single mutants since neither of them showed steady state current. But this is expected if the two-tier barrier is acting as a steric hindrance in series as is observed in the pore radius measurement where two layers of hydrophobic barriers blocks the pore at the bottom of S6 and the permeation pathway could only be widened by reducing both residues forming the barrier.

**Figure 2:**
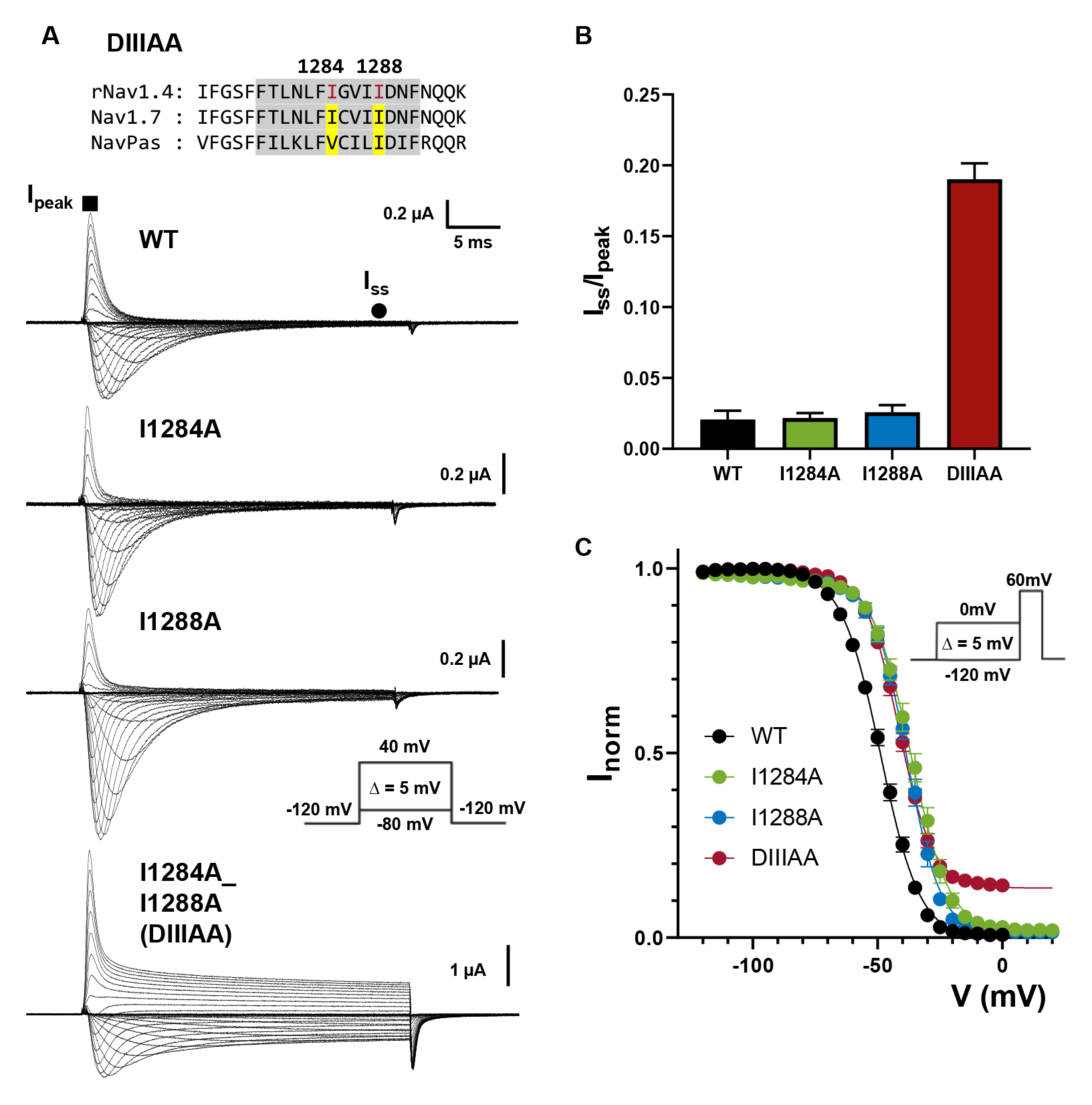
Double alanine mutation in DIII S6 relieves fast inactivation on rNav1.4. A) Top: sequence alignment of DIII S6 showing identified residues (I1284 and I1288, highlighted in yellow). Bottom: Representative ionic traces for the wild-type (WT), single (I1284A and I1288A) and double alanine (I1284A_I1288A, named DIIIAA) mutations of the identified residues on rNav1.4. The ionic conditions used was 57.5 Na^+^ outside and 12 Na^+^ inside. Inset shows the voltage protocol. B) Ratio of steady state current (I_ss_, ●) at the end of depolarization of 30ms over the peak current (I_peak_, ▪) taken at +60mV. C) Voltage-dependence of inactivation (h-infinity curve) for WT (black), I1284A (green), I1288A (blue) and DIIIAA (red). The curves were made by plotting the normalized peak ionic current (Inorm) at the test pulse (60mV) after 50ms conditioning prepulse (voltage protocol in inset). The lines represent the fit to a two-state model (Equation 1). Data are shown as Mean ± SEM. Number of cells tested are in table I.

To characterize the conformational changes along the fast inactivation pathway, we measured gating charge immobilization in DIIIAA. As Nav channels enter fast inactivation, VSDs in DIII and DIV get trapped in the up conformation and become immobilized. As a result, upon hyperpolarization, VSDs in DIII and DIV move after the channels transitioned out of fast inactivated state, manifesting as a slow component in the off-gating current (16). Therefore, the amount of immobilized charge, and the time course of immobilization provide a direct measurement of conformational changes along the fast inactivation pathway, independent of ionic current. In WT, almost all channels are inactivated a few milliseconds after depolarization (Fig. 2A). Consistent with this, a second slow kinetic component develops in the off-gating current after milliseconds of depolarization (Fig. 3A). Despite the presence of steady state ionic current (Figure 2A), a similar pattern of charge immobilization was observed in the off-gating currents of DIIIAA (Fig. 3B). In WT and DIIIAA about 60% of the charge was immobilized and both constructs shared a similar time course (Fig. 3C). This result suggests that DIIIAA channels transits completely into the fast inactivated state and the steady current observed cannot be explained assuming a subpopulation of channels unable to enter the inactivated state. The fact that the time course of immobilization is about the same between WT and DIIIAA, indicates that the rate constants of transitioning in and out of fast inactivation were not significantly affected by the double alanine mutation. This is further supported by the essentially identical kinetics of the ionic current inactivation in DIIIAA and WT (Fig. 3D). Thus, the kinetic pathway from open state to fast inactivated state was not dramatically altered in DIIIAA, consistent with the charge immobilization results.

**Figure 3:**
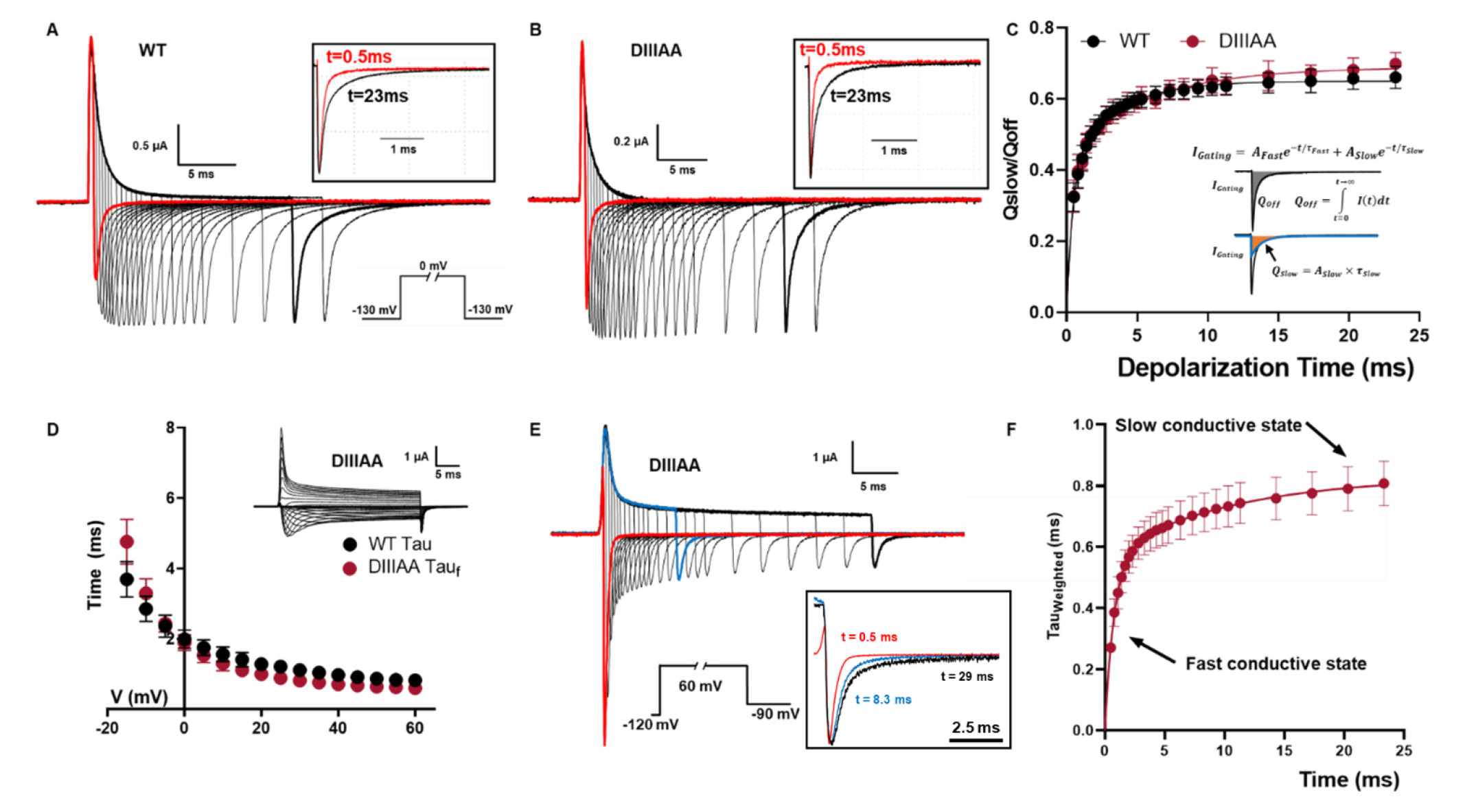
Evidence of a conductive “inactivated” state. Representative traces of gating current used for charge immobilization measurements for WT (A) and DIIIAA (B), bottom inset in (A) shows the voltage protocol. Black square top inset shows the comparison of the normalized gating charge for a 0.5 ms and 23 ms depolarization pulse. C) Fraction of immobilized charge vs depolarization time for WT (black) and DIIIAA mutant (red). The inset illustrates the method used for determination of the off-gating components using exponential fitting (Detailed in methods). D) Fast Inactivation time constant for WT (black) and DIIIAA (red). Fast inactivation kinetics were characterized by fitting a single exponential function for WT and two-exponential function for DIIIAA. Inset shows representative DIIIAA current traces in response to a voltage protocol shown in Figure2A. E) DIIIAA currents in response to depolarized voltage step to +60mV with variable duration followed by a hyperpolarized voltage step to −90mV. The inset shows the normalized tail currents at 3 depolarization times. F) Weighted tail currents time constant vs depolarization time for DIIIAA mutant (Details in methods). Data are shown as Mean ± SEM.

Upon short voltage depolarization, tail currents from DIIIAA mutant were fast and exhibited a single time constant of ∼ 0.3 ms. But with longer depolarizations, the tails slowed down considerably (Fig 3. E). This was evidenced by the appearance of a second slow component reaching a time constant close to 0.8ms, pointing to the existence of two distinct conductive states (Fig 3.F). The change in tails kinetics followed the time course of the fast inactivation in DIIIAA (Supplementary Figure 3). Overall, it is apparent that the double alanine mutation in DIII S6 created a “leaky” inactivated state where all the Nav channels could still transition into the fast inactivated state through a pathway similar to that in WT, as suggested by the charge immobilization and the kinetics results. However, the final inactivated state is still conductive, manifested as a second conductive state.

We find no evidence to suggest that the reduction in the side chain volume in DIIIAA has any effect on the “leakiness” of the closed state at rest which suggests that the high energy barrier associated to the sharp reduction in conductance in the closed state must be linked to a separate set of S6 residues.

### The “leaky” inactivated state is less selective for Na^+^ ion

In certain voltage ranges, we noticed that the ionic currents went from inward to outward during the pulse in the DIIIA mutant. This was shown as a fast inward and a steady-state outward component (Fig. 4 A and Supplementary Figure 4 A). Since both current components are blocked by TTX (Figure 4 B, Supplementary Figure 4 D) it suggests that both are permeating through the pore and that Nav selectivity might be changing throughout the voltage pulse. Indeed, the reversal potential for the peak current, which is related to the first component, is significantly more positive than to the one for the steady state current, related to the second component (Fig. 4 A). When external Na^+^ was increased from 57.5 to 90 mM both components were shifted to more depolarized voltages: from 35 to 46 mV for the peak component and from 18 to 30 mV for the steady state component (Fig 4 A, B). When only K^+^ is used as permeant ion there is no crossing in the ionic currents and the reversal potential for the peak and steady state are the same (Figure 4 C). Once K^+^ was exchanged by Na^+^ in the external solution, the crossing reappears, showing that this process is reversible (Supplementary Figure 5). Therefore, our results uncovered a surprising change in the ionic selectivity of DIIIAA during the time course of depolarization. To further explore this change in selectivity, we measured the selectivity at different time points during a depolarizing voltage pulse (60 mV). This was done by measuring the instantaneous I-V curve at different times during depolarization, computing the reversal potential as a function of time, and using the GHK equation (Equation 6) as a framework to analyze the data (Figure 4D, E, Supplementary Figure 6, and Methods). Under bi-ionic conditions we can calculate the relative permeability between Na^+^ and K^+^ ions (P_Na_/P_K_) from the reversal potential measurements. At the beginning of the pulse, the selectivity of DIIIAA towards Na^+^ was high, similar to the WT (P_Na_/P_K_ = 11) but almost three time less selective later in the pulse (P_Na_/P_K_ = 4) (Fig. 4F). The time course of the permeability change followed the time course of the fast inactivation, indicating that the conformational changes associated with fast inactivation might also trigger the change in selectivity (Fig. 4 G). These results point to the existence of an allosteric coupling pathway between S6 and the SF. Independent of the underlying coupling mechanism, these results show, unequivocally, that the bulky hydrophobic residues identified form part of the inactivation gate.

**Figure 4:**
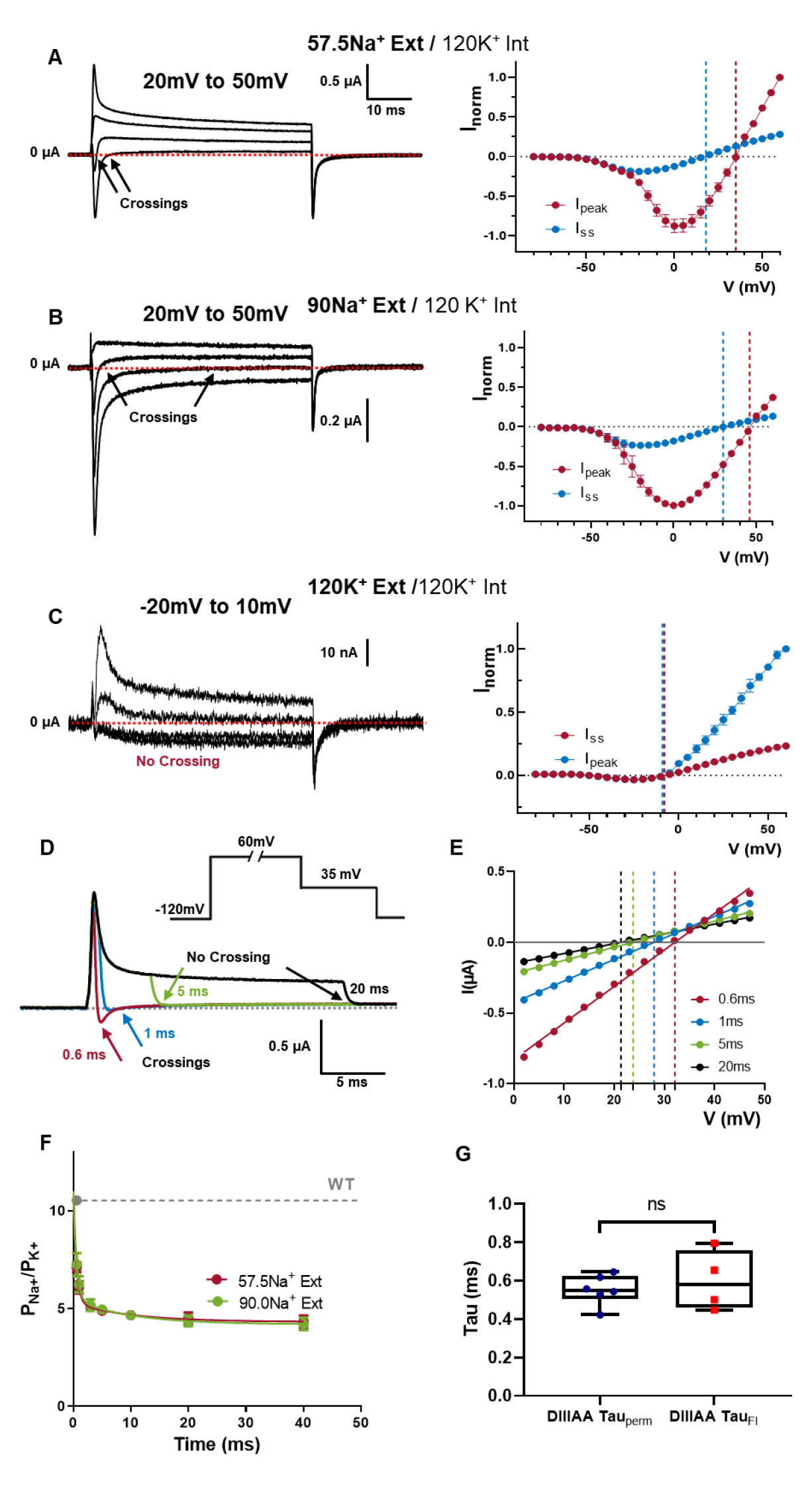
Time dependent selectivity changes in DIIIAA. DIIIAA depolarization activated currents at different voltages (range shown on top) using 57.5 mM external Na^+^ (A), 90 mM external Na^+^ (B) or 120 external K^+^ (C), on the right I-V curves for the peak (red) and steady state (blue) current. The reversal potential for each component is denoted with a corresponding dashed line. D) Example of the changes in direction for the tail currents. E) Instantaneous I-V curves after a 0.6 (red) 1 (blue) 5 (green) and 20 (black) milliseconds depolarization times. The solid lines represent a linear fit used to obtain the reversal potential (indicated by vertical dashed lines). F) Relative Na/K permeability vs time for DIIIAA under different ionic conditions (red, 57.5 mM Na external and green, 90 mM Na external concentration. Internal solution 120 mM K. Dashed line shows the WT permeability and solid lines represent an exponential fit to obtain the time constant. G) Fast time constants for the change in selectivity (DIVAA Tau_perm_) and fast inactivation (DIIIAA Tau_FI_).

### “Leaky” inactivated state can be prevented by removing fast inactivation in DIIIAA

One interesting feature of the present results is that, at hyperpolarized voltages (from - 55mV to −30mV), the DIIIAA mutant displays slower ionic currents kinetics but no apparent inactivation, a phenomenon not observed for the single mutants and WT (Supplementary Fig. 7 A-D). When compared to WT Nav, the voltage dependence of activation (G-V curve) of DIIIAA was shifted 8.5mV to the left (calculated at the steady-state), whereas the two single alanine mutants were right shifted (15.2mV for I1284A; 9.6mV for I1288A) (Fig. 5A – Supplementary Table 2). Despite this, the voltage dependence of the voltage sensor movement (Q-V curve) showed a shift to the right in DIIIAA (8.5mV) when compared to the WT (Fig 5. B, C – Supplementary Table 3). This suggests a possible increase in the coupling between the VSD and PD for DIIIAA. However, one simple explanation for this paradoxical result was that the “leaky” inactivated state present in DIIIAA shows ionic currents at less depolarized voltages, therefore, shifting the G-V curves to the left, since Nav inactivation precedes activation in voltage (35, 37), this would explain the early component in the current as flowing through the leaky inactivated state.

**Figure 5:**
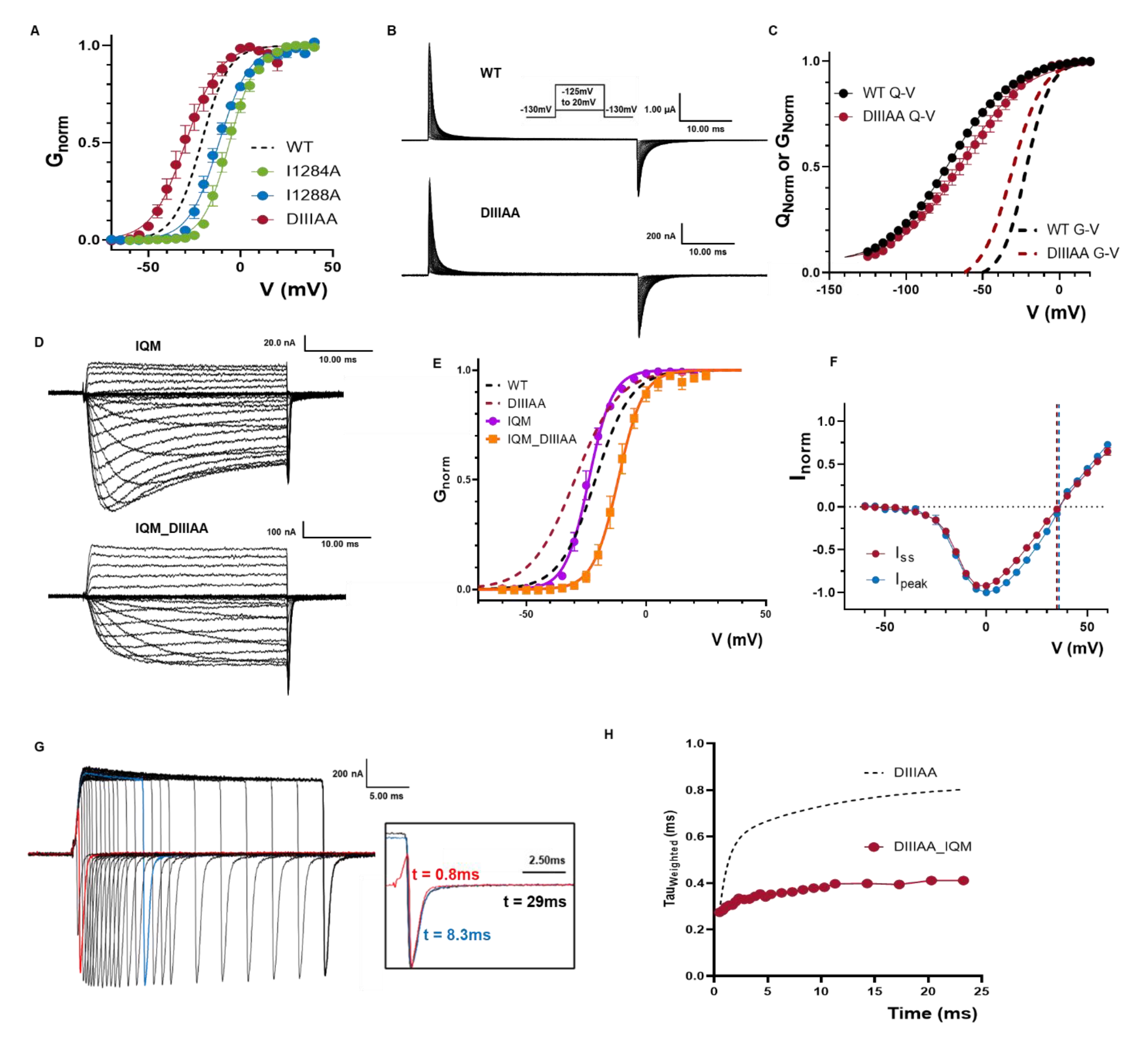
Preventing IFM binding avoids the DIIIAA phenotypes. A) G-V curves for WT (black dashed line), I1284A (green), I1288A (blue) and DIIIAA (red). GV curves were calculated from the peak for WT, I1284A and I1288A whereas for DIIIAA it was calculated from the steady state currents. B) Representative gating current traces for WT and DIIIAA. Inset is the voltage protocol. C) Q-V curves for WT (black) and DIIIAA (red). The dashed lines indicate the corresponding G-V curves. D) Representative ionic current traces for IQM and IQM_DIIIAA. E) G-V curves for IQM (magenta) and IQM_DIIIAA (orange). Dashed lines show the WT and DIIIAA for comparison. F) IQM_DIIIAA IV curves for the peak (red) and steady state (blue) current. The reversal potential for each component is denoted with a corresponding dashed line. G) IQM_DIIIAA currents in response to depolarized voltage step to +60mV with variable duration followed by a hyperpolarized voltage step to −90mV. The inset shows the normalized tail currents at 3 depolarization times. H) Weighted tail currents time constant vs depolarization time for IQM_DIIIAA (red points) compared to DIIIAA (dashed line).

In order to test our hypothesis, we introduced the F1304Q mutation in the IFM motif in both WT (IQM) and DIIIAA (IQM_DIIIAA) constructs to remove fast inactivation. Within the previously accepted “hinged-lid” model, IQM was supposed to remove fast inactivation by preventing the binding of IFM to its ultimate receptor in the open pore. In the current framework where we demonstrate that the residues at S6 forms the fast inactivation gate, we interpret the IFM motif, instead of being the final effector, acts more likely as a transducer that couples the VSD movement in DIV which triggers the fast inactivation, to the pore where the inactivation gate ultimately closes. By mutating F1304 to Q, the energetic barrier becomes so high that effectively, the conformational changes associated with fast inactivation is terminated before reaching the pore. By preventing fast inactivation, we expect to remove the features associated with the leaky inactivated state, namely: the slow component on the tail currents, the change in selectivity and the shift in the G-V curve. In WT Nav, IQM removes most of the fast inactivation without significant shifts in the G-V curve (Fig. 5 D, E, Supplementary Table 2). In contrast, in IQM_DIIIAA, the G-V curve was shifted to the right (9.6 mV) compared to the WT, yet we saw no changes in selectivity and tail currents did not change with the depolarization time (Fig. 5 D-H). Since the IQM mutation itself does not change the G-V curve and removes inactivation, our results indicate that: i) the “leaky” inactivated state appears downstream of IFM binding; ii) the apparent shift seen in DIIIAA G-V curve was likely due to the channel entering into the “leaky” fast inactivation state from a closed state which contributed to the early current seen at hyperpolarized voltages; and iii) the change in selectivity becomes apparent as channels transition into the fast inactivated state.

### DIV S6 is part of the inactivation gate

DIV, similar to DIII, has been shown to have specialized roles in fast inactivation. We identified two large residues at the bottom of DIV S6 based on the sequence alignment, I1587 and L1591 (Fig. 6 A). When I1587 and L1591 residues were mutated to alanine simultaneously (DIVAA), significant steady-state current was observed across all voltages tested (around 13% at 60mV) (Fig. 6A, B). However, single alanine mutations (I1587A, L1591A) did not drastically change the voltage dependence of fast inactivation (Fig. 6C – Supplementary Table 1). Despite the steady state current in DIVAA, the amount of charge immobilization is similar to the WT and DIIIAA (Fig. 6 D, E). Furthermore, in DIVAA, the fast inactivation process had similar time course as the WT and DIIIAA. These results show that DIVAA channels can also transit into the fast inactivated state but, similarly to DIIIAA, the fast inactivation gate is partially open creating a “leaky” inactivated state. Consistently, two components of current with different reversal potentials are observed in DIVAA (Figure 6F and Supplementary Figure 8). After determining the relative permeability between Na^+^ and K^+^ as a function of time, we observed a decrease in selectivity in DIVAA. At the onset of the pulse, the selectivity of DIVAA towards Na^+^ was high (P_Na_/ P_K_=13), but later in the pulse the relative permeability for Na decreased (P_Na_/P_K_=2) (Fig. 6G). The change in selectivity follows the kinetics of inactivation as evidenced by their similar time constants (Fig. 6H). The gating charge movement and the activation process are not drastically affected by the double alanine mutations DIVAA (Supplementary Figure 9).

**Figure 6:**
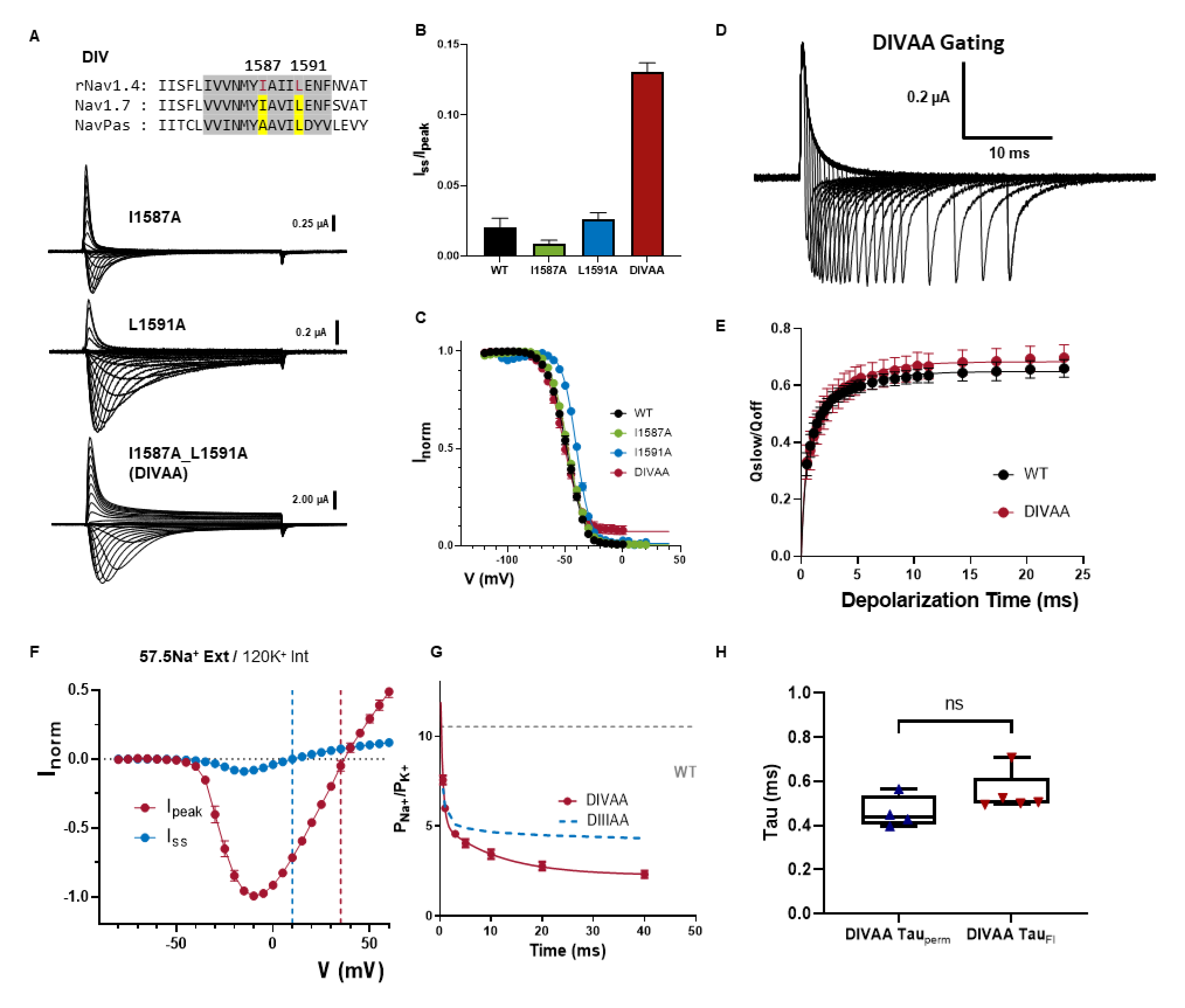
Alanine mutations in DIV also produced a leaky inactivated state. A) Top: sequencing alignment of DIV S6 showing identified residues (I1587 and L1591, highlighted in yellow). Bottom: Representative ionic traces for single (I1587A and L1591A) and double alanine (I1587A_L1591A – DIVAA) mutations. The ionic conditions used was 57.5 Na^+^ outside and 12 Na^+^ inside. B) Ratio of steady state current (I_ss_) at the end of depolarization over the peak current (I_peak_) taken at +60mV. C) Voltage-dependence of inactivation (h- infinity curve) for WT (black), I1587A (green), L1591A (blue) and DIVAA (red). D) Representative gating currents traces for charge immobilization measurements for DIVAA. E) Fraction of immobilized charge vs depolarization time for WT (black) and DIVAA mutant (red). F) IV curves for the peak (red) and steady state (blue) current for DIVAA. The reversal potential for each component is denoted with a corresponding dashed line. F) Relative Na/K permeability vs time for DIVAA (57.5 mM Na external and 120 mM K internal). Dashed grey and blue lines show the WT and DIIIIAA permeability, respectively. Solid line represents an exponential fit to obtain the time constant. G) Fast time constants for the change in selectivity (DIVAA Tau_perm_) and fast inactivation (DIVAA Tau_FI_). Data are shown as Mean ± SEM.

Based on the many parallels between DIIIAA and DIVAA, we believed that the effects produced by the double alanine mutation in DIII, and DIV share the same underlying mechanism and the identified hydrophobic residues in DIII together with the ones in DIV are part of the fast inactivation gate. To further demonstrate the shared role of DIII and DIV as part of the inactivation gate, we mutated all four residues identified to alanine (DIII_IVAA). The effect of the quadruple alanine mutation was drastic. This mutant exacerbated the effects seen in DIIIAA and DIVAA alone. Firstly, most fast inactivation was removed (Fig. 7 A and Supplementary 10). while the kinetics of the residual fast inactivation remained comparable to the WT. Secondly, the ionic selectivity became severely impaired and time dependent as evidenced by the fast inward and steady-state outward component at some voltages (Fig.7A-B). The P_Na_/P_K_ at the onset of the pulse was already ∼2.4 and at steady-state 1.5 (Fig. 7C). These results demonstrate that all four residues are part of the fast inactivation gate and reinforce the hypothesis of a coupling between the bottom of S6 region and the SF

**Figure 7:**
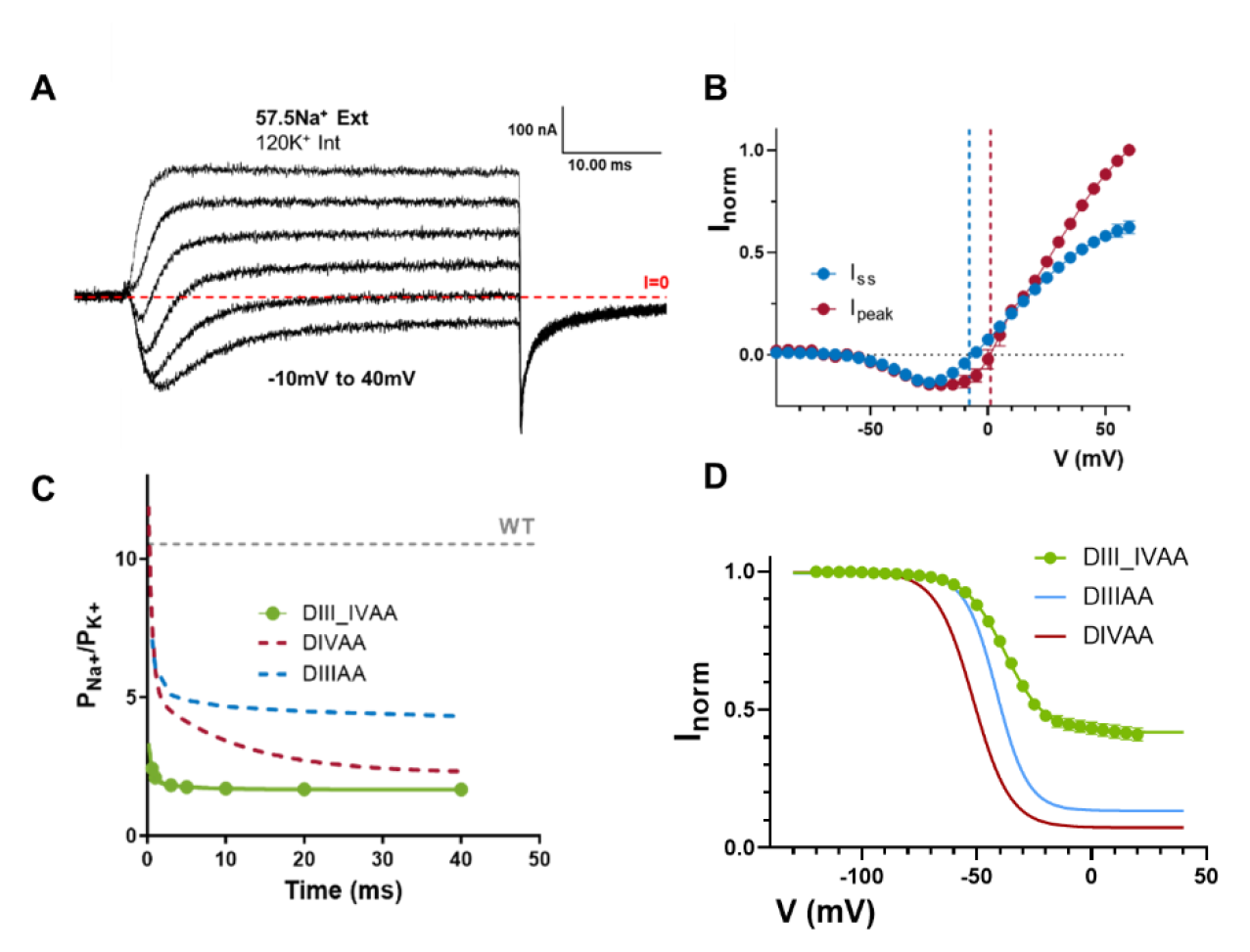
Quadruple alanine mutation of the identified residues in DIII & DIV S6. A) Representative current traces elicited at different voltages (−10 to 40 mV) for the quadruple alanine mutation (I1284A_I1288A_I1587A_L1591A, DIIIAA_IVAA) using 57.5 mM external Na and 120 internal K. (B) IV curves for the peak (red) and steady state (blue) current. The reversal potential for each component is denoted with a corresponding dashed line. C) Relative Na/K permeability vs time for DIVAA (57.5 mM Na external and 120 mM K internal). Dashed grey, blue and red lines show the WT, DIIIIAA and DIVAA permeability, respectively. Solid line represents an exponential fit to obtain the time constant. D) h-infinity curve for DIII_IVAA, DIIIAA and DIVAA. Significant fast inactivation was removed in DIII_DIVAA.

### Effects in DI and DII are not consistent with participation in the inactivation gate

The effects of alanine mutations in DI and DII were drastically different from DIII and DIV. In DI S6 out of the two identified residues, only one residue was large and hydrophobic (L437), and the second identified residue was an alanine (A441). Since residue 441 is already an alanine we only tested a single alanine mutation (L437, DIA). In DIA, a small amount of steady state current was detected as well as a ∼15 mV right-shifted in the voltage dependence of activation (Figure 8 A-B). Despite this effect on steady state current, the h-infinity curve shows that channels can fully inactivate (Figure 8C). Furthermore, there is no evidence of a time dependent change in the selectivity, as evidenced by the lack of two components in the ionic currents (Supplementary figure 12). On the other hand, when we performed the double alanine mutation on the identified residues in DII S6 (L792A_L796A, DIIAA), the effects were different to what was described for DI, DIII and DIV. In DIIAA, the ionic current showed robust and complete fast inactivation across all voltages tested (Figure 8D). Despite the lack of steady state current, at the end of the depolarizing pulse a large tail current was observed. The most likely explanation for the origin of these “tail” currents is that they are gating currents. This correlates with the observation of two distinct current components during the depolarizing pulses. One component that is fast and always outward and another that follows the reversal potential for sodium and is similar to the ionic currents of the WT channels (Figure 8 E). The tail currents are observed after the ionic currents are completely inactivated and its appearance correlates with the development of the first component indicating that indeed during the depolarizing pulse, the first component is gating current, and the second component is ionic current (Figure 8E, Supplementary figure 11). Since the ratio between the peak ionic and gating currents in WT sodium channels is in the order of 50 to 1, and in DIIAA is in the order of 2 to 1, these results indicate that there exists significant uncoupling between PD and VSD in DIIAA. Altogether, the results of the DIA and DIIAA despite their possible relevance do not show unequivocally that these residues form part of the inactivation gate, unlike DIIIAA and DIVAA.

**Figure 8:**
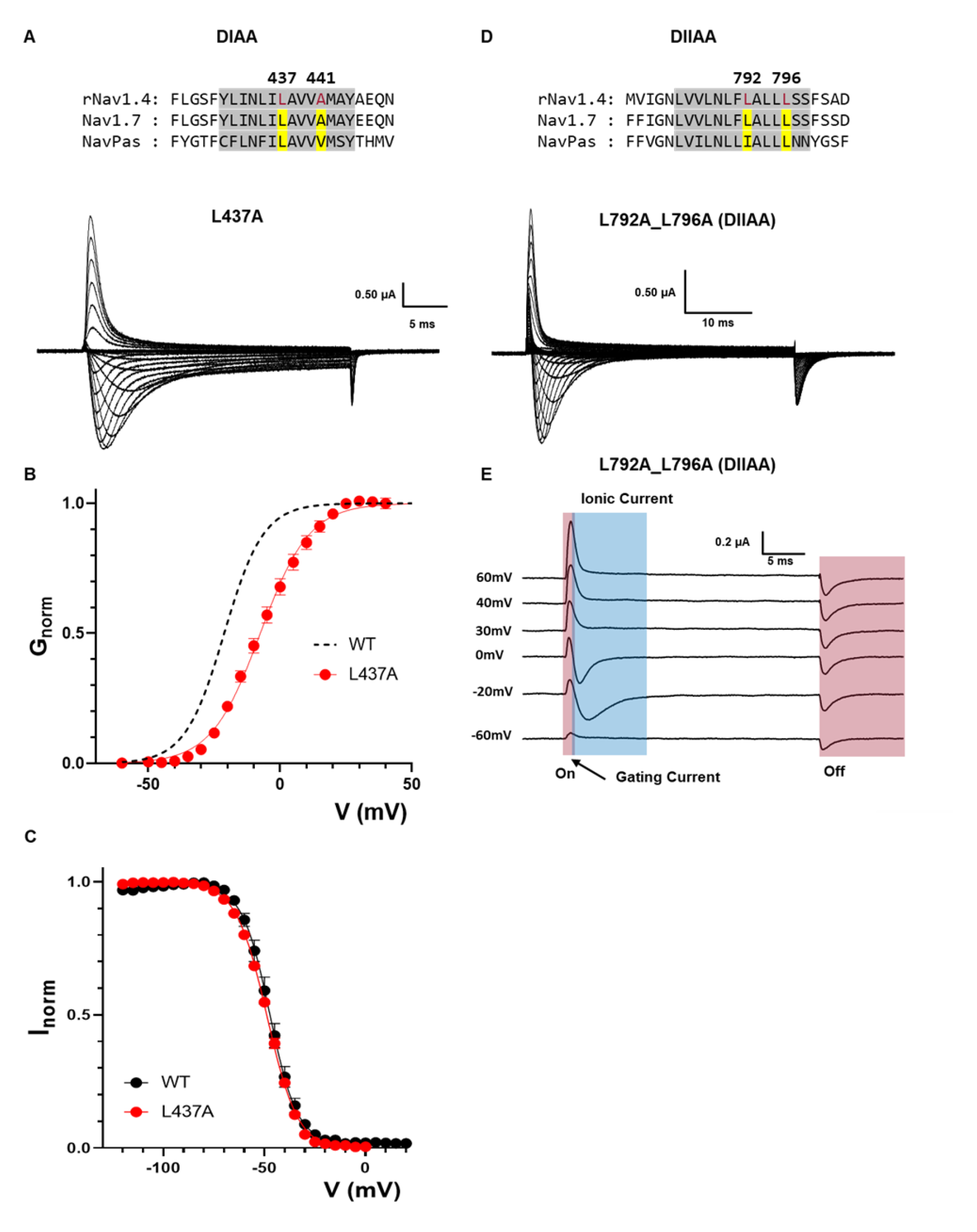
Effects of alanine mutations in DI and DII S6. A) Top: sequencing alignment of DI S6 showing identified residues (L437 and A441, highlighted in yellow). Bottom: Representative ionic traces for L437A. The ionic conditions used was 57.5 Na^+^ outside and 12 Na^+^ inside. B) G-V curves for WT (black dashed line) and L437A (red). GV curves were calculated from the peak. C) H-infinity curve for WT (black) and L437A (red). D) Top: sequencing alignment of DII S6 showing identified residues (L792 and L796, highlighted in yellow). Bottom: Representative current traces for L792A_L796A (DIIAA). The ionic conditions used was 57.5 Na^+^ outside and 12 Na^+^ inside. E) Detail of DIIAA the gating and ionic current components. The red and blue shadowing indicate the presence of gating and ionic currents respectively.

## Discussion

### The controversies on fast inactivation gate: location and identity

Fast inactivation is one of the defining features in Nav channels and has implications in many physiological processes. The IFM motif has long been considered the inactivation particle that blocks the pore during a “ball and chain”-type fast inactivation (18, 21). Manipulations to the IFM motif can lead to severe impairments in fast inactivation, for instance, by mutating IFM to QQQ, fast inactivation can be removed completely (18, 38). Moreover, adding a small peptide that contains the IFM sequence into the intracellular side of the membrane can partially rescue the fast inactivation in the QQQ mutant, which suggests IFM could block directly the open pore (39). However, when the first structure was solved for mammalian Nav channels(22), it clearly contradicted the *status quo* model. Several Nav structures showed, not only that the IFM motif was in principle far away from the pore in the putative inactivated state, but also that some residues previously identified to influence fast inactivation did not interact directly with IFM at all, L437 and A438 being the prime example (40).

An important “missing link” in the Nav fast inactivation saga, relates to the fact that, while there are multiple high resolution Nav cryoEM structures, actually assigning a precise functional state to individual structures is among the current major challenges in structural biology. While it is possible that the structures were resolved in the fast inactivated state, it is also possible that they may have been captured in a slow inactivated state, a different process from fast inactivation (37, 41) with putatively significantly different structural features. Yet, if we assume that some of the existing structures represent a fully or partially fast inactivated conformation, the S6 helices and associated bulky hydrophobic side chains logically emerge as strong candidates to block ion flow during fast inactivation (Figure 1).

It is worth pointing out that the different Nav structures do not offer a clearcut consensus regarding the exact residues that could form the fast inactivation gate. Different S6 residues have been proposed based on the different structures (Nav1.5 (33) and Nav1.7(23, 28)). However, detergents were found lodging at the bundle crossing region of many Navs, potentially distorting the conformation of the channel around that region and further clouding potential interpretations. Here, our experiments and interpretations are based on two structures that are free from this problem, NavPas and Nav1.7 M11. Even though NavPas has never been functionally characterized and Nav1.7 M11 has 11 stabilizing mutations incorporated, our functional data support the idea that the pore profiles from these two channels are likely to be similar to the ones under physiological conditions and represent the fast inactivated pore. Therefore, they serve as a framework to analyze the fast inactivation gate.

In DIII and DIV, none of the single alanine mutation yielded significant change to the fast inactivation. Only when both residues in the same domain or when quadruple residues were mutated to alanine, did we start to see significant amount of steady state current from a leaky inactivated state (Figures 2,4,6,7). This result clearly demonstrates that an S6-based fast inactivation comprises at least two layers of residues with side chains pointing into the pore, making the hydrophobic region longer along the pore. The requirement of two layers to make the gate might also be the reason why many previous single alanine scanning studies on S6 did not show a clear effect and therefore were not able to identify the fast inactivation gate (42): the two rings of identified bulky hydrophobic residues act *in series*.

Our results show that in DIIIAA and DIVAA, the channels go through all the conformational changes along the fast inactivation pathway. This is supported by several lines of evidence: I) the kinetics of the residual fast inactivation are similar to the WT (Figure 3) II) gating charge immobilization, a process closely linked to fast inactivation, occurs in DIIIAA and DIVAA at the same level and with the same kinetics as the WT (Figure 3 and 6) and III) the slowing down of the tail currents and the change in selectivity, associated with the leaky inactivated state, have the same time course as inactivation (Figure 3, 4 and 6). Finally, removal of the fast inactivation process in the IQM mutant relieves the effects observed in DIIIAA (Figure 5). The most direct interpretation of these results is that the double alanine mutations DIIIAA and DIVAA did not affect the sequence of conformational changes that lead to inactivation but rather make the inactivated state conductive.

### Potential coupling between DIII, DIV S6 helices and the SF

An unexpected result from our work was that the ion selectivity changed concomitantly following the kinetics of fast inactivation in DIIIAA and DIVAA (Figures 4 and 6). Na^+^ ions were more permeable than K^+^ ions during the early component (before fast inactivation set in) while the relative K^+^ permeability increased over time. Given that TTX blocks both components of the current completely, Na^+^ and K^+^ are clearly conducted through the pore domain of the channel (Figure 3). It is also worth noting that the general behavior of DIIIAA and DIVAA show a striking resemblance with that of batrachotoxin (BTX) modified Nav channels. BTX modifies Nav channels in three major ways: i) it removes fast inactivation; ii) it shifts the GV curve to the left; iii) it alters the selectivity of the channel (43). Parallels could be found in DIIIAA and DIVAA corresponding to all three modifications. More intriguingly, BTX is hypothesized to bind to the neurotoxin receptor site 2 which is close to the mutated residues (I433, N434, L437 in DI S6 and F1579 N1584 in DIV S6 have been shown to be crucial for BTX binding) (44). Therefore, it is likely that they share a similar underlying mechanism, so that BTX evolved as a toxin to disrupt the inactivation gate at S6 even more effectively than DIIIAA and DIVAA.

Due to the asymmetrical assembly of Nav channels, each domain is only similar, not identical to the others, each one playing slightly different roles in the activation and inactivation processes (34, 45). In this work, we also observed the effects of this asymmetry. In DI, only one of the identified residues was bulky and hydrophobic (L437) while the second position was an alanine (A441). L437A produces slight changes to fast inactivation (Figure 8). The apparent conclusion would be that either the residues at DI S6 did not contribute enough to the fast inactivation or our analysis of the structures was not able to identify the correct residues in DI for fast inactivation gate. However, we cannot rule out the importance of residue L437 since previous work has demonstrated the double L437C/A438W mutant (CW) removes close to 90% of Nav fast inactivation (40). Our data show that residues located at DI S6 are able to influence fast inactivation, however, the effects observed are not sufficient to unequivocally assign it as part of the fast inactivation gate.

The DII S6, double alanine (DIIAA) seems to not be involved in the inactivation gate. We demonstrated that in DIIAA the gating current became disproportionally larger compared to the ionic current (Figure 8), therefore, it is likely that the double alanine mutation in DII S6 decreases the open probability of the channel. DII VSD has been shown to be involved mostly in channel activation instead of fast inactivation(45). We suspect that this apparent decrease in P_O_ was a result of an uncoupling between the VSD and the PD during activation. Our results suggest that the bottom of S6 region in DII plays a different role from the other domains. Even though, our data support an uncoupling between VSD and PD, we cannot rule out the possibility of a drastic enhancement of inactivation similar to the effects of W434F mutation in Shaker K^+^ channel (46, 47). Therefore, it requires further investigation to determine the underlying mechanism of this uncoupling.

### Fast inactivation as a multistep process

Our results show that the bottom of the S6 region of the pore serves as the fast inactivation gate. One implication of these results is that fast inactivation is a multistep process and the IFM motif binding is only one step, albeit critical, in a whole sequence of conformational changes. Previously, the fast inactivation process was largely described as a two-step process: VSD in DIV activates, exposing the binding pocket for the IFM motif and IFM motif binds, blocking the pore. Our experiments provide a different interpretation of the mechanism of fast inactivation in Nav channels as a series of conformational changes. In this new model (Figure 9), the activation of VSD in DIV triggers the movement of the IFM motif by exposing a binding pocket. The IFM motif then binds to its binding pocket that is away from pore. The binding site of the IFM is far from the S6 regions, therefore it is expected that the binding event of the IFM is transduced to the S6 gate in the pore through a yet to be defined pathway. Once the movement is allosterically relayed to the S6 segments, the large residues at the bottom of S6 occlude the pore and stop the permeation. The exact nature of these conformational changes and the residues involved in the third step is still elusive and the underlying mechanism is yet to be determined. However, either a rotation, translocation, or a slippage of the S6 helices towards the pore could lead to the pore closure during fast inactivation.

**Figure 9:**
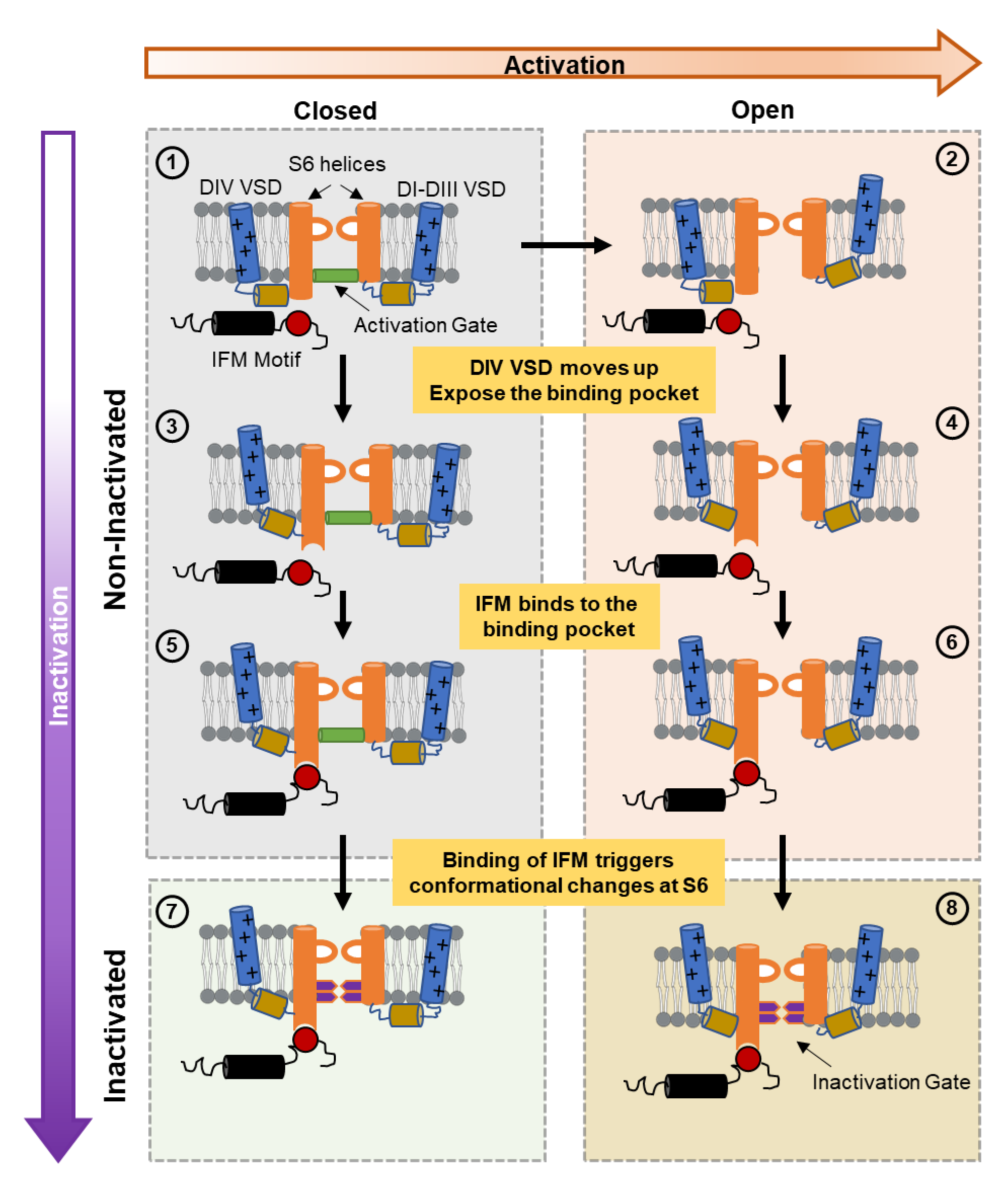
A general model for fast inactivation from open or closed states. For simplicity, the multiple-step activation is shown as a single transition between the last closed state and the open state in the horizontal direction (rightward) and vertical transitions represent the inactivation path (downward). ① represents the Nav channel in the last closed state. ② represents the channel with the VSD of DI-DIII in the active (up) position, DIV VSD is in the resting (down) position and a conducting pore: an open channel. Transitions from ① to ③ and ② to ④ occur when DIV VSD moves up, exposing the binding pocket for the IFM motif outside the pore region. Once the binding pocket becomes available, the IFM motif binds and transitions from ③ to ⑤ and ④ to ⑥ occur. The binding of the IFM motif triggers conformational changes that are conducted to the pore region closing the two-layered fast inactivation gate, leading to the closed inactivated state in ⑦ or the open inactivated state in ⑧.

Finally, our results show a change in selectivity during the leaky inactivated state which implies a previously unrecognized communication between S6 and the selectivity filter. This interaction opens the intriguing possibility that during the normal inactivated state the sodium channel is less selective to Na^+^. Regardless, it is clear that in light of the current work, many previously identified mutations that influence fast inactivation might require reinterpretation.

## Acknowledgement

We would like to thank Dr. Eduardo Perozo for his thoughtful comments and careful editing of the manuscript; Dr. Rong Shen for his help on calculating the pore radius in the cryo-EM structures.

This work is supported by National Institutes of Health Award R01GM030376 and National Science Foundation Award QuBBE QLCI (NSF OMA-2121044)

## Material and Method

### Structure analysis and pore radius calculation

Deposits of structures and electron density maps were analyzed using ChimeraX (48)and VMD (49). The pore radius was calculated by HOLE (31). Sequencing alignment was performed using SnapGene.

### Site-Directed Mutagenesis and RNA synthesis

Rat Nav1.4 α and β1 subunit cloned into pBSTA vector, flanked by β-globin sequences was used (ref of the original clones). Mutagenesis was performed utilizing mismatch mutagenesis primers in a two-staged PCR reaction for best efficiency. The PCR products were used to transform *E. coli* XL-gold competent cells. After ampicillin resistance screening plasmid were purified using standard DNA miniprep (macherey-nagel). Each DNA was sequenced to confirm the desired mutation and absence of off target mutations.

DNAs were linearized at the unique NotI restriction site and then transcribed into complementary RNA using T7 *in-vitro* transcription kits (ambion).

### Channel Expression in *Xenopus* Oocytes

Ovaries of *Xenopus laevis* were purchased from XENOPUS1. The follicular membrane was digested by collagenase 2 mg/ml supplemented with bovine serum albumin (BSA) 1mg/ml. After defolliculation, Stage V-VI oocytes were microinjected with 50-150 ng of premixed cRNA with 1:1 molar ratio of α and β1 subunits. Injected oocytes were incubated at 18°C for 1-5 days prior to recording in SOS solution (in mM: 100 NaCl, 5 KCl, 2 CaCl_2_, 0.1 EDTA and 10 HEPES at pH 7.4) supplemented with 50 µg/mL gentamycin.

### Electrophysiology

Ionic and gating currents were recorded using the cut-open voltage-clamp technique (50). Micropipettes with resistance between 0.3 and 0.8 MΩ were used to measure the internal voltage of the oocytes. Current data were filtered at 20 kHz with a low-pass Bessel filter online and subsequently sampled by a 16-bit analog-to-digital converter at 1 MHz. Transient capacitive currents were first compensated by a dedicated circuit and then minimized by an online P/N protocol (51). Temperature was maintained at 11.5 ± 1 °C by a feedback Peltier device throughout the experiment. To lower the series resistance, 1M cesium methylsulfonate (MES) solution was used in the shortened agar bridges. For ionic current experiments, external solutions consisted of in mM: X NaMES, 120-X N-methyl-D-glucamine (NMG) MES, 2 CaMES, 10 HEPES and 0.1 EDTA, pH = 7.4; Sodium internal solutions consisted of in mM: 12 NaMES, 108NMG MES, 10 HEPES and 2 EGTA, pH =7.4; Potassium internal solution consisted of in mM: 120 KMES, 10 HEPES and 2 EGTA, pH = 7.4. Subtraction voltage was set between −80 mV to −120mV with a P/-4 protocol for ionic currents. Gating currents were recorded in 120 NMG MES, 2CaMES, 10 HEPES, 0.1 EDTA external solution and 120 NMG MES, 10 HEPES, 2EGTA internal solution (in mM). Both solutions were titrated to pH 7.40. To block ionic conductance, 750nM of tetrodotoxin (TTX) was added into the external bath solution. Subtraction voltage was set at +20mV with a P/4/ protocol. For records containing both ionic and gating currents, the recording solutions were consistent with the solutions used for ionic current experiment. To minimize the loss of gating currents due to P/-4 protocol while maintaining good resolution of ionic currents and good health of the oocytes, −130mV was chosen to be the subtraction voltage. Voltage clamp speed, measured by capacitive transients, yielded a time constant around 75µs and the kinetics of gating current was reliably resolved.

### Internal solution dialysis for oocytes

To ensure the ionic composition of the oocyte cytoplasm we exchanged the internal solution in oocytes by incubation in divalent free solution made up of in mM: 120 KMES, 10 HEPES, 0.1 EDTA and 2EGTA, pH = 7.4. The oocytes were incubated in the dialysis solution between 30 min to 1hour at room temperature prior to experiments.

### Data Analysis

GraphPad prism, Excel, Matlab and in-house software (Analysis and GPatchM) was used to process all the results.

I) Steady-state fast inactivation curve (h infinity curve) was assembled by plotting normalized peak Na^+^ current during test pulse against the conditioning pulse voltage and fitted using a two-state Boltzmann model,

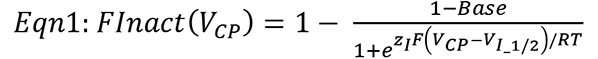

Where Finact (V_CP_) is the normalized remaining current after given conditioning pulse at voltage V_CP_. Base, z_I_ and V_I_1/2_ are the percent of non-inactivating current, the apparent charge of the process and the half inactivation voltage, respectively. R, T and F represent typical values for the gas constant, experiment temperature and Faraday constant respectively.

<u>II</u>) To determine the percentage of charge immobilization, off-gating current was fitted two-component exponential decay.

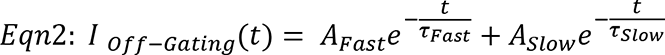

Where A_Fast_, τ_Fast_, A_Slow_ and τ_Slow_ represent amplitudes and time constants of the fast and slow components respectively.

The amount of charge in the slow component was calculated by multiplying A_Slow_ and τ_Slow_. Percentage of charge immobilization was then determined by dividing the amount of charge in the slow component by the total amount of off-gating charge, as is shown:

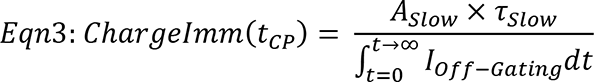

Where ChargeImm(t_CP_) is the percentage of charge immobilization after a conditioning pulse of duration t_CP_ and 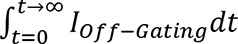 Is the total gating charge as measured by the integral of the off-gating current the current.

III) The time constant of inactivation or deactivation kinetics were calculated by fitting the ionic currents with either one or two exponential decays using the general equation:

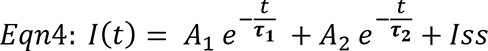

Where A_1_, τ_1_, A_2_ and τ_2_ represent amplitudes and time constants of the first and second components, respectively. I_SS_ is the steady state current. When one exponential was used, the term of the second exponential component was eliminated.

To obtain the weighted time constant we used the following relationship:

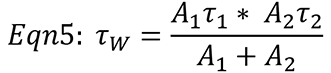

IV) Relative permeability between Na^+^ and K^+^ ions was determined in bionic environmental with 120mM intracellular K^+^ and different extracellular Na^+^ concentration. The reversal potential was determined by fitting the instantaneous IV curve with a linear relationship and locating the intersection point on the voltage axis. The reversal potentials as a function of depolarization time were subsequently used to determine the relative permeability using Equation 6, derived from GHK equation (52, 53).

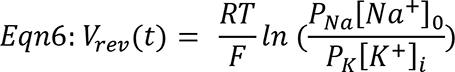

where V_rev_(t) is the reversal potential after depolarizing pulse of duration t; P_Na_ and P_K_ represent relative permeability of Na^+^ and K^+^ respectively; [Na^+^]_o_ and [K^+^]_I_ represent extracellular Na^+^ concentration and intracellular K^+^ concentration, respectively.

<u>V</u>) The ionic conductance (G(V_m_)) was calculated by dividing either the peak current or steady state current by the experimentally determined driving force, at each depolarizing voltage (V_m_). Subsequently, the curve was normalized to the maximal conductance across all voltages (Gmax) to obtain the conductance versus voltage (G-V) curves. The G-V curves were fitted using a two-state model:

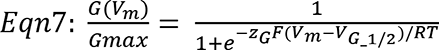

Where V_G_1/2_ is the half activation voltage and z_G_ is the apparent charge.

<u>VI)</u> Total gating charges movement during voltage pulses was measured by integrating the gating currents. Charge-voltage (Q-V) curves were obtained by plotting the normalized total charge movement (Q_Norm_) at each depolarizing voltage and fitted using a two-state model (Equation 8),

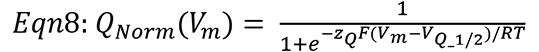

Where V_Q_1/2_ is the gating half activation voltage and z_Q_ is the apparent charge.

## Author contributions

Performed and analyzed experiments: YL

Interpreted results: YL, CB, BP FB

Conceptualization: YL, CB, BP, FB

Writing: YL, CB, BP, FB

Supervision: FB

## Data and materials availability

All data presented in the paper will be made available to readers upon request.

