## Supplementary Figures for "A Mechanistic Reinterpretation of Fast Inactivation in Voltage-Gated Na^+^ Channels"

### Supplementary Information

Supplementary Table 1: Fit parameters of the voltage-dependence of inactivation for all the channels tested.

| | $V_{I\_1/2}$ | $z_I$ | Base | n |
|---|---|---|---|---|
| **WT** | **-48.83±0.18** | **-3.08** | **/** | **8** |
| **I1284A** | **-37.10±0.32** | **-2.96** | **0.01±0.0069** | **6** |
| **I1288A** | **-38.82±0.27** | **-3.37** | **0.01±0.0060** | **6** |
| **DIIIAA** | **-41.52±0.26** | **-3.51** | **0.13±0.0064** | **4** |
| **I1587A** | **-47.58±0.13** | **-3.22** | **0.00±0.0026** | **5** |
| **I1591A** | **-40.34±0.20** | **-4.04** | **0.01±0.0046** | **5** |
| **DIVAA** | **-51.62±0.32** | **-3.02** | **0.07±0.0069** | **8** |

Supplementary Table 2: Fit parameters of the voltage-dependence of activation for all the channels tested.

| | $V_{G\_1/2}$ | $z_G$ | n |
|---|---|---|---|
| **WT** | **-21.19±0.45** | **3.30** | **4** |
| **I1284A** | **-5.951±0.33** | **3.42** | **4** |
| **I1288A** | **-11.62±0.32** | **2.97** | **6** |
| **DIIIAA** | **-29.72±0.62** | **2.67** | **4** |
| **IQM** | **-23.89±0.26** | **4.97** | **4** |
| **IQM_DIIIAA** | **-11.65±0.38** | **4.68** | **4** |
| **I1587A** | **-12.48±0.31** | **3.33** | **5** |

| | | | |
|---|---|---|---|
| **I1591A** | **-18.31±0.27** | **2.75** | **5** |
| **DIVAA** | **-21.97±0.60** | **4.05** | **6** |
| **L437A** | **-7.511±0.30** | **2.64** | **8** |

Supplementary Table 3: Fit parameters of the voltage-dependence of charge movement.

| | $V_{Q\_1/2}$ | $z_Q$ | **n** |
|---|---|---|---|
| **WT** | **-74.12±0.28** | **1.16** | **6** |
| **DIIIAA** | **-66.34±0.61** | **1.12** | **6** |
| **DIVAA** | **-73.13±0.23** | **1.07** | **7** |

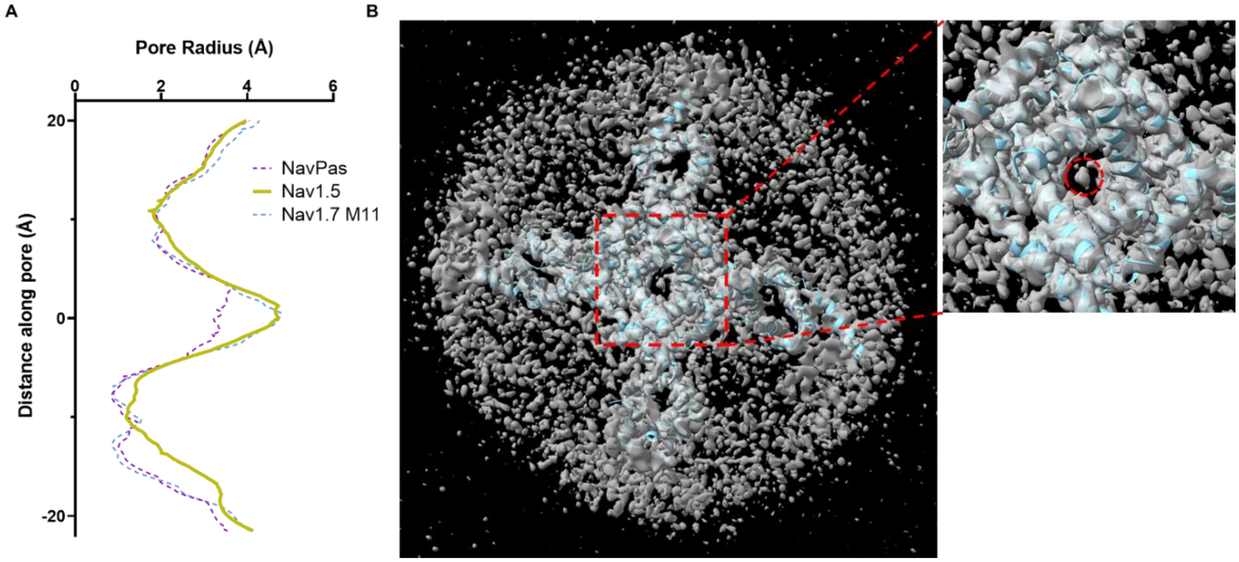


**Supplementary Figure 1**. Differences in pore radius between NavPas, Nav1.7M11 and Nav1.5 (6UZ3). A) The narrowest part in NavPas and Nav1.7 M11 structure is much broader than the Nav1.5 structure. B) Intracellular view of the electron density map overlayed with the structure model. There exists unidentified density inserting into the pore (highlighted in red circle).

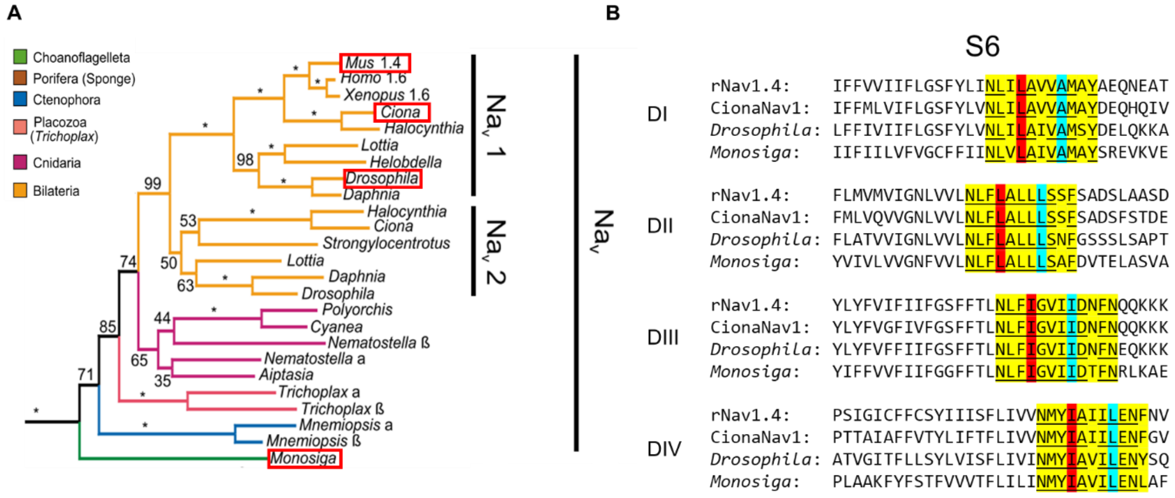

**Supplementary Figure 2**. Sequence alignment of Nav from different species. The identified residues are very conserved across many different species. A) is adapted from Liebeskind et al., 2011(54).

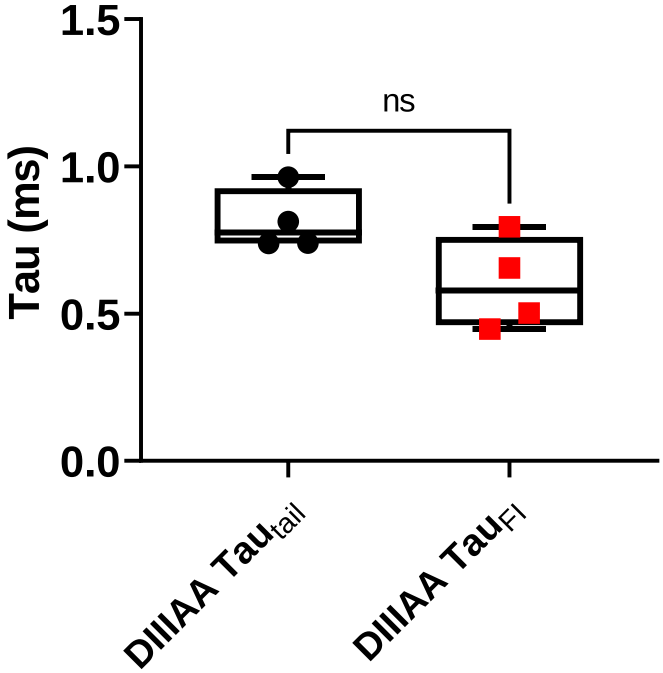


**Supplementary Figure 3**. Fat time constant of the slowing down of the tail current and fast inactivation. No significant difference was found (unpaired t-test with Welch's correction, $p > 0.5$)

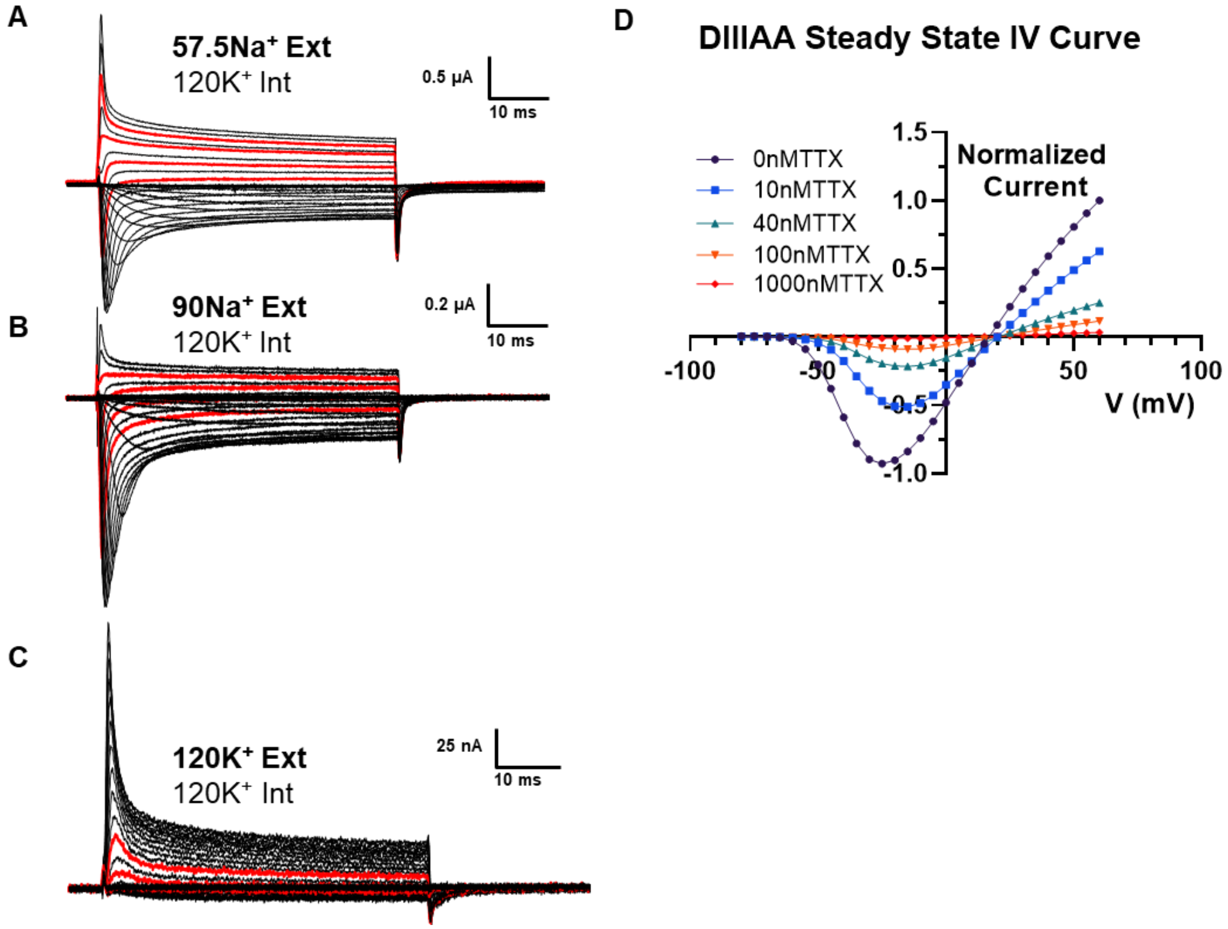


**Supplementary Figure 4**. DIIIAA ionic currents elicited by a set of voltage pulses (from -80 to + 60 mV every 5 mV), using 57.5 mM external Na (A), 90 mM external Na (B) or 120 external K (C). The red lines indicate the traces used in figure 4 (D). TTX could block the current efficiently

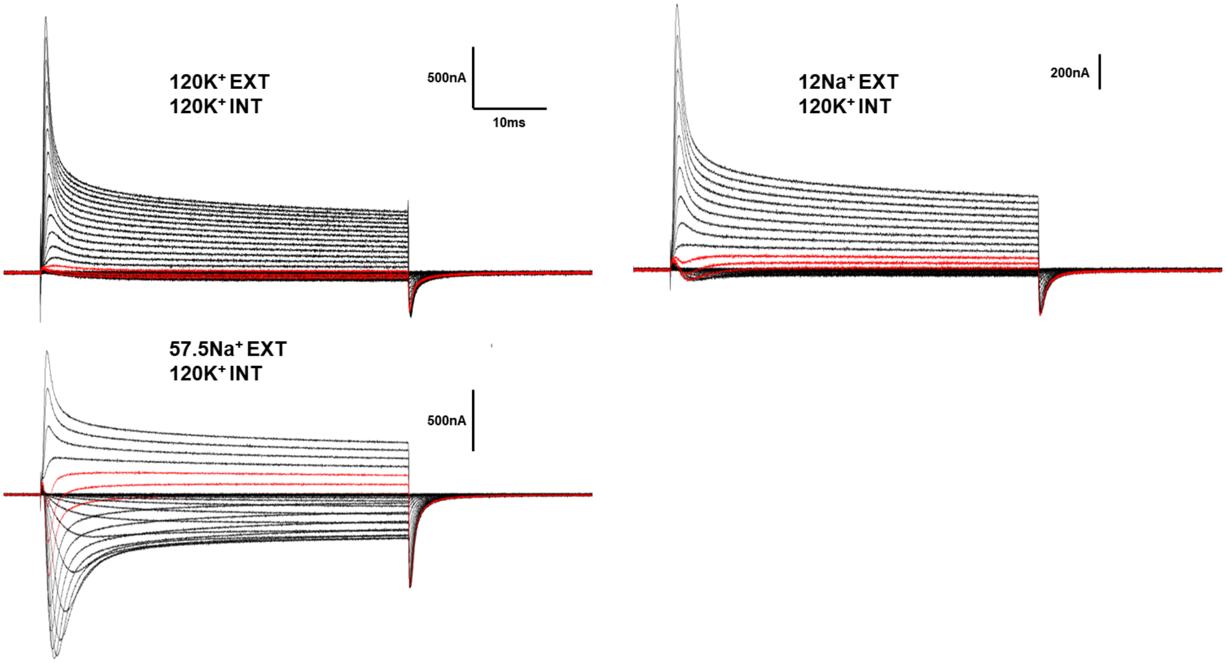


**Supplementary Figure 5.** Once $K^+$ was exchanged by $Na^+$ in the external solution, the crossing reappears, showing the reversibility of the two components of current. The experiment shown above was performed on the same oocyte starting with external $120K^+$ then to $12Na^+$ and finally in $57.5Na^+$. The traces near the reversible potential are highlighted in red.

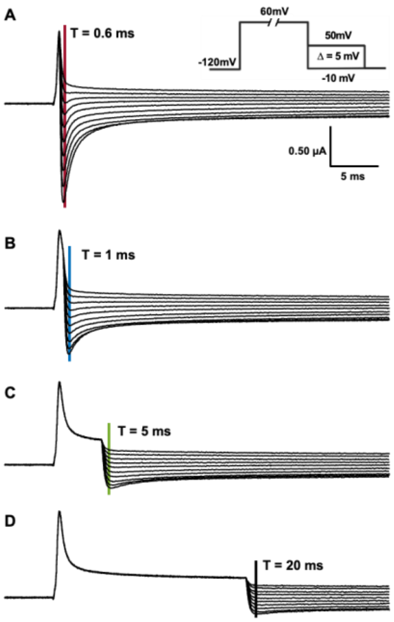


**Supplementary Figure 6.** Representative current traces used to obtain the instantaneous IV curves and calculate their reversal potential at 0.6 (A), 1 (B), 5 (C) and 20 (D) ms depolarization time. The vertical lines are the isochrones used to calculate the instantaneous IV. Voltage protocol shown at inset in A.

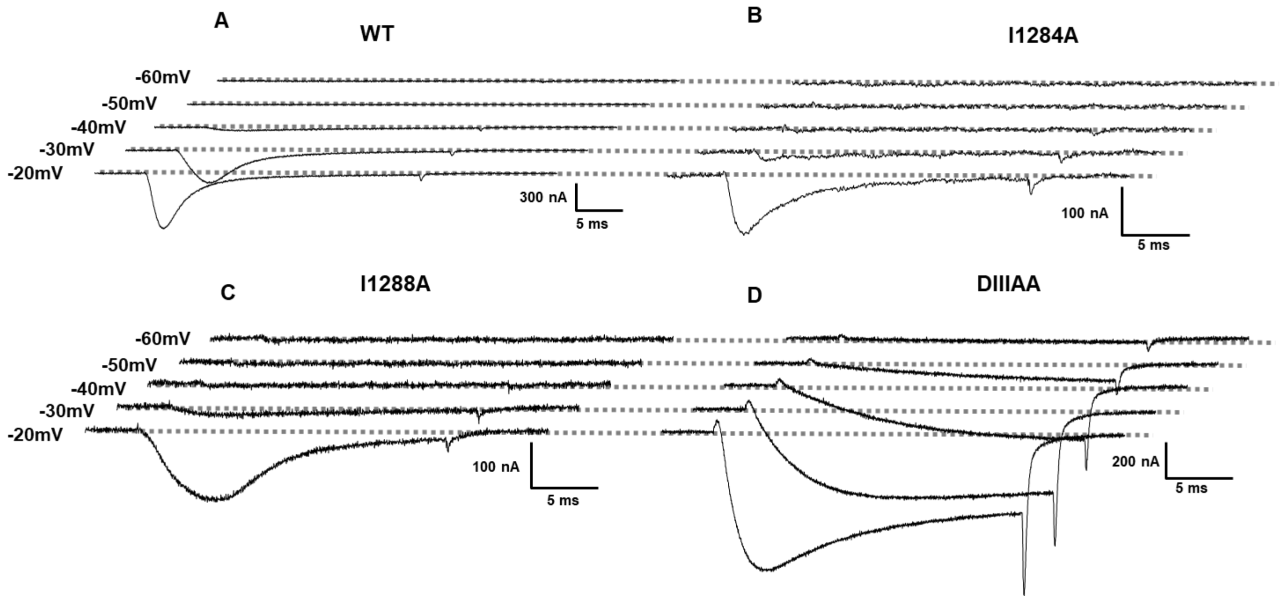


**Supplementary Figure 7:** Ionic currents at hyperpolarized voltages in the WT (A), I284A (B), I288A (C) and DIIIAA mutant (D). The dashed lines indicate the zero current baseline.

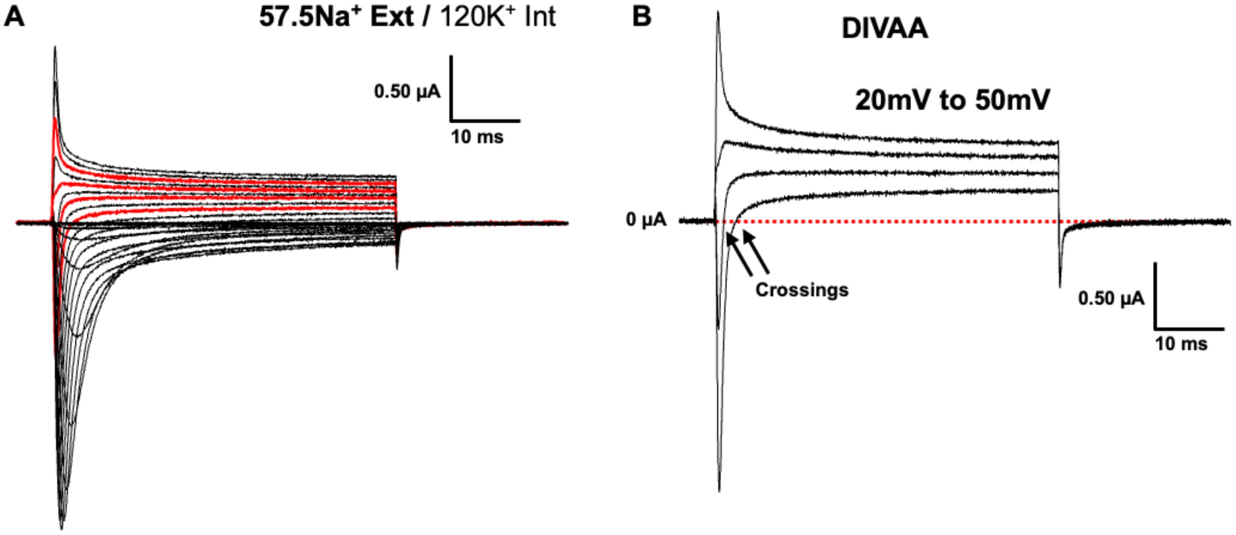


**Supplementary Figure 8.** DIVAA ionic currents elicited by a set of voltage pulses (from -80 to + 60 mV every 5 mV), using 57.5 mM external Na. (A). B) For clarity purposes we show the red current traces from A. Arrows indicate the crossing points.

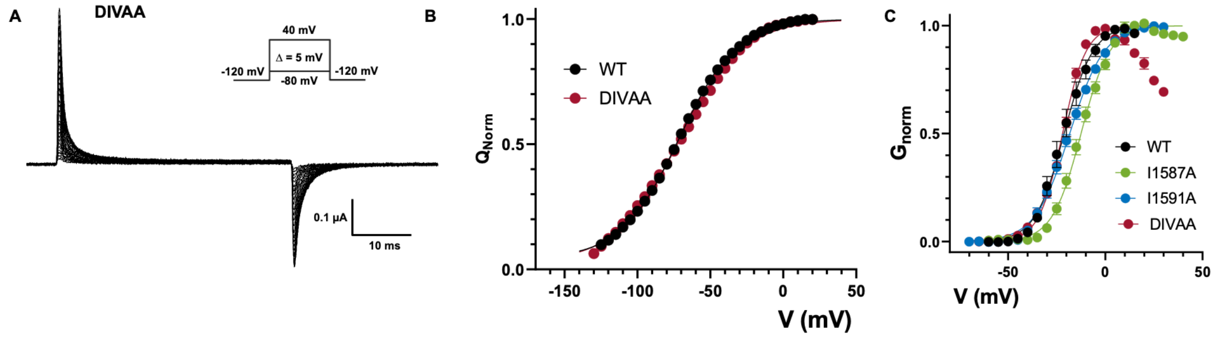


**Supplementary Figure 9**: Gating properties of alanine mutations at DIV S6. A) Representative gating current traces for DIVAA. Inset is the voltage protocol. B) Q-V curves for WT (black) and DIVAA (red). C) G-V curves for WT (black), I1587A (green), L1591A (blue) and DIVAA (red). G-V curves were calculated from the peak for WT, I1587A and L1591A whereas for DIVAA it was calculated from the steady state currents.

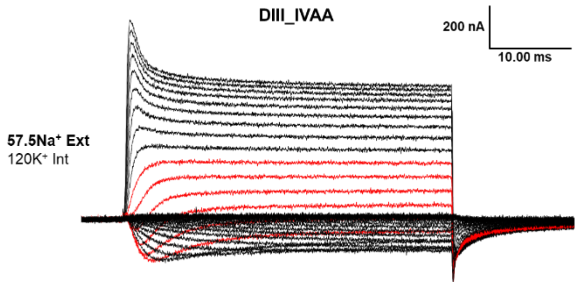


**Supplementary Figure 10**: Quadruple alanine mutation of the identified residues in DIII & DIV S6. Depolarization activated currents for the quadruple alanine mutation (I1284A_I1288A_I1587A_L1591A, DIIIAA_IVAA) at different voltages (using 57.5 mM external Na and 120 internal K

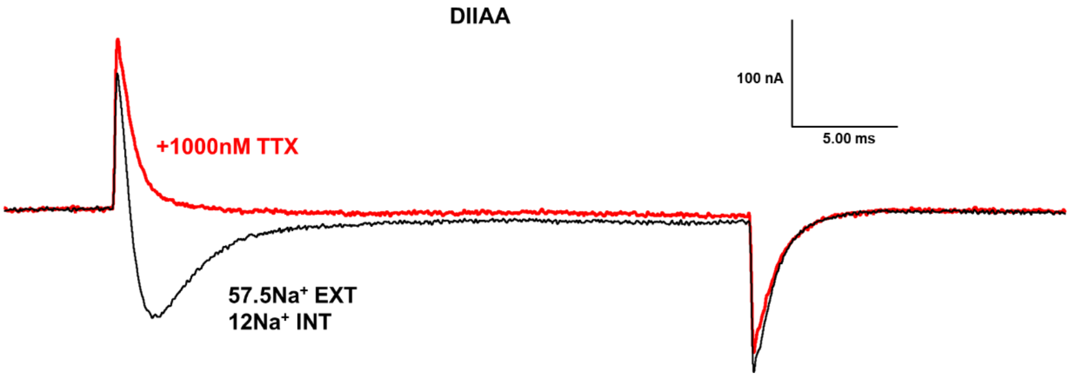


**Supplementary Figure 11:** DIIAA has disproportionally large gating current. Shown here is the current elicited at 0mV without (black trace) and with (red) TTX. Clearly the early component and off-tail current originate from gating current.

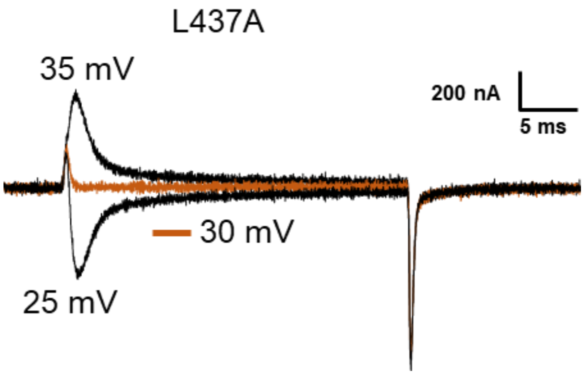


**Supplementary Figure 12:** Lack of two components on DIA ionic currents. Representative ionic current traces for DIA at 3 voltages close to the reversal potential (~30 mV).
